# The primary and secondary immune response to Epstein-Barr virus infection of human tonsil organoids

**DOI:** 10.64898/2026.01.16.698390

**Authors:** Álvaro F. García-Jiménez, Luis Ignacio González-Granado, Andrea Sánchez de la Cruz, Ignacio Jiménez-Huerta, Mar Valés-Gómez, Hugh T. Reyburn

## Abstract

Epstein-Barr virus (EBV) is a human herpesvirus that causes acute infectious mononucleosis (IM) and is associated with cancer and autoimmune disease. Humans are the only natural host for EBV and humanised mice are the only small animal model of infection. Consequently, although IM in adults following primary infection has been studied intensively, little is known about the virological and immunological events that occur during the initial phases of the virus-host interaction. EBV is usually transmitted via saliva exchange, and is thought to infect its main host, the human B cell, in mucosal secondary lymphoid tissues, such as the tonsils. Thus, to address this gap in knowledge, we have studied the immune response to EBV infection using tonsil organoids as a model that allows us to address the greater systemic complexity present at the site of infection in the oropharynx. EBV infection is efficiently controlled when the tonsils are derived from “virus-experienced” individuals and CD8^+^ memory T lymphocytes expressing CD103 play a leading role in this immune response. In contrast, in primary infections, immune control of lymphoblastoid cell proliferation is much less effective, and a key factor restricting the immune response is the secretion of the immunomodulatory viral IL-10 molecule. These results highlight the importance of the host cytotoxic response at the site of infection and demonstrate that immune evasion molecules appear to be crucial for EBV-infected B cells to elude the local immune response and so disseminate the infection throughout the organism.

## INTRODUCTION

Epstein-Barr virus (EBV) has co-evolved with humans for millennia and is a highly successful pathogen, infecting around 95% of people worldwide^1,2^. Infection with this gamma-herpesvirus normally occurs in young children and acquisition of the virus in childhood establishes a lifelong asymptomatic infection^1,2^. However, this virus-host balance depends critically on effective immune surveillance and if this immunity is disturbed any one of a range of virus-associated diseases can occur. For example, the reactivation of EBV on immunosuppression is a major cause of morbidity and mortality after transplantation^3^. EBV is also associated with around 2% of all human cancers (some 300,000 new cases and 200,000 cancer deaths every year)^4,5^. Finally, compelling epidemiological data implicate EBV infection as a key necessary step for the development of multiple sclerosis (MS)^6^, the most prevalent autoimmune disease of the central nervous system, and systemic lupus erythematosus (SLE)^7,8^.

Immune control of EBV is thought to depend mainly on EBV-specific T-cell responses that are readily detected in healthy carriers^9^. Studies of patients with inborn errors of immunity where chronic EBV viremia and/or EBV-associated disease are common clinical features confirm the importance of EBV-specific CD8 T cells and NK cells for immune control of EBV-infected B cells^10^. However, such analyses do not provide a complete picture of overall immune control, because they generally depend on sampling blood as a readily accessible source of immune cells. Specifically, the role of virus-specific tissue resident lymphocytes in immunity to EBV is poorly understood. It has been shown that in long-term virus carriers, EBV-specific CD8 T cells are enriched in tonsil compared with blood^11,12^, but evidently these data were obtained long past the primary stages of the interaction of the virus with the immune system.

Increased knowledge of the immunological events in asymptomatic primary infection would provide a better understanding of the key elements of protective immunity against EBV. However, for obvious reasons, we are almost completely ignorant about the virological and immunological events that occur in asymptomatic primary infection, especially in the earliest phases of the virus-host interaction. Most work studying the development of T cell responses during primary infection has focussed on people identified as recently infected with EBV through the overt symptoms of infectious mononucleosis. Although valuable, there are some important limitations to these studies. First, IM represents an immunopathological condition, where patient symptoms arise from the excessive CD8 T lymphocyte responses^9^. Thus, whether IM is a useful model for immune control of primary EBV infection or simply a consequence of inefficient immunity is unclear. Second, viral infection occurs several weeks prior to symptoms developing and samples being taken^13,14^. Therefore, our knowledge about the initial stages of EBV infection and the immune response to that viral challenge in humans is, at best, fragmentary.

Since the initial phases of EBV infection occur in the oropharynx where tonsils are located^1,2,9^, we have studied the very first phases of virus infection and the immune response to that infection using a novel, physiologically relevant model: EBV infection of tonsil organoids. In this manuscript, we show that the activation of tissue resident CD8 T cell populations distinguishes seropositive from seronegative individuals who cannot control primary EBV infection. We also identify the production of the IL-10 homologue encoded by EBV as an important factor limiting the development of T cell-mediated immunity early in primary infection. These data provide a deeper understanding of the immune response to EBV at the likely site of infection and highlight the role of immunoevasins in the very earliest phases of the virus-host interaction.

## RESULTS

### 1. Cytotoxic lymphocytes are activated after EBV infection of tonsil organoids

Inspired by the tonsil organoid system developed by Wagar *et al*^15^, an EBV infection model was established using tonsil samples from pediatric tonsillectomy (**Supplementary Figure 1A**). After analysis of samples from multiple donors, two outcomes of infection were observed. In cultures that developed a secondary response (IgM-IgG+ antibodies in supernatant, seropositive donor), the proliferation of virus-infected cells was efficiently controlled. Whereas in cultures with features of a primary response (IgM+IgG- antibodies detected in culture, seronegative donor) infection led to a significant expansion of lymphoblastoid cells (CD19^+^CD38^high^) (**Figure 1A-C, Supplementary Figure 1B,C**). These cells could be confirmed as EBV infected when GFP-expressing EBV was used (**Supplementary Figure 1D**). No correlation was found between tonsil donor age and either IgG or IgM antibody responses (**Supplementary Figure 1E**). These observations led to two questions: what components of the immune system mediate the efficient control of EBV replication in secondary responses? Secondly, what factors might be modulated to enhance the primary immune response to EBV?

**Figure 1.**
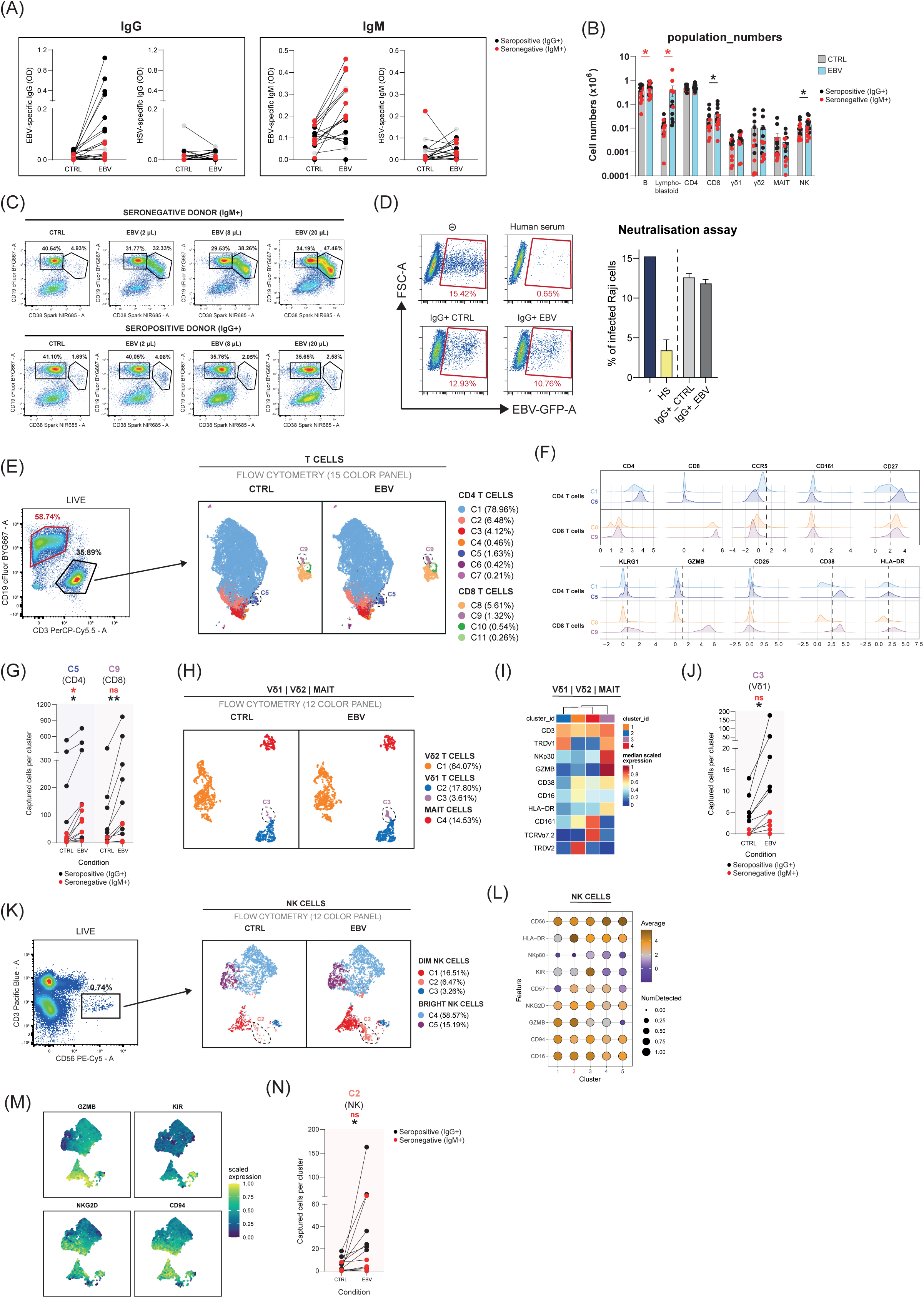
Activation of cytotoxic cells in tonsils of IgG+ individuals after infection with EBV. (A) ELISAs quantifying IgM and IgG antibodies specific for EBV or HSV in the supernatant of tonsil organoids 10 days after infection with EBV. (B) Number of cells in control and infected conditions at day 10. Paired sample t-test (*<0.05). (C) Examples of lymphoblastoid cell detection (CD19^+^CD38^high^) after titration of EBV in IgM+ and IgG+ individuals at day 10. (D) Neutralising activity of human sera (n=3, 1:100) and IgG+ tonsil supernatants, either non-infected or infected (n=7, 1:2), against EBV-GFP infection of Raji cells. (E) UMAP of T cells split by condition. 10,000 cells per donor were plotted. The most differential clusters (C5 and C9) were highlighted. (F) Histograms representing the expression of markers in C1, C5, C8, and C9 in the flow cytometry T cell panel. (G) Quantification of cells in C5 and C9 in control and infected conditions at day 10. Paired sample t-test (*<0.05, **<0.01). (H) UMAP of Vδ1 | Vδ2 | MAIT split by condition. All cells were plotted. The most differential cluster (C3) was highlighted. (I) Heatmap representing the expresion of markers in the clusters of the flow cytometry Vδ1 | Vδ2 | MAIT panel. (J) Quantification of cells in C3 in control and infected conditions at day 10. Paired sample t-test (*<0.05). (K) UMAP of NK cells split by condition. All cells were plotted. The most differential cluster (C2) was highlighted. (L) Dot-plot representing the expression of markers in the clusters of the flow cytometry NK cell panel. (M) Feature-plots highlighting the expression of some markers in NK cells. (N) Quantification of cells in C2 in control and infected conditions at day 10. Paired sample t-test (*<0.05).

As little significant production of virus-neutralising antibodies was detected in tonsil cultures that controlled the infection (**Figure 1D**), we decided to characterise the cellular immune response against EBV. For this purpose, flow cytometry panels were optimised based on prior data from our group studying PBMC infection models^16^ (**Supplementary Figure 1F**). In IgG+ individuals, a greater number of CD8 T lymphocytes (CD27^+^KLRG1^+^GZMB^+^) were activated compared to CD4 T lymphocytes (**Figure 1E-G, Supplementary Figure 1G,H**), even though only low numbers of CD8 T cells are present in tonsils. Interestingly, in some donors, a population of cytotoxic CD4 T cells with high expression of GZMB and KLRG1 was also detected (**Supplementary Figure 1I,J)**, while regulatory T cells (FOXP3^+^) were only captured in low proportions (**Supplementary Figure 1K**). Other populations of cytotoxic lymphocytes, such as γδ T cells expressing the Vδ1 TCR that upregulated NKp30 (**Figure 1H-J, Supplementary Figure 1L**), and NK cells with high expression of NKG2D and CD94 (**Figure 1K-N, Supplementary Figure 1M,N**), were also detected in these samples. However, the extent of activation of all these cell types was much lower in IgM+ individuals.

No obvious signs of activation of innate lymphoid cells (ILCs), predominantly ILC1 (CRTH2^-^Kit^-^) and ILC3 (CRTH2^-^Kit^+^NKp44^+^) were noted after EBV infection, at least as judged by HLA-DR expression (**Supplementary Figure 2A-C**). In contrast, macrophages, which were mainly of the CD14^+^CD16^+^ phenotype, increased in number and showed some upregulation of the activation marker CD80 (**Supplementary Figure 2D-F**).

### <u>2.</u> Activation of EBV-specific CD103^+^ CD8 T cells correlates with efficient control of EBV infection

Since the ability to control virus infection in this model was a feature of virus-experienced donors, we used spectral cytometry (**Supplementary Figure 3A)** to analyse how the different human T cell memory populations found in tonsils responded to EBV. Significantly increased numbers of memory CD8 T cells after EBV infection were only noted in those individuals who made anti-EBV IgG antibody responses *in vitro* (**Supplementary Figure 3B**). Detailed analysis of these data showed that these individuals were able to increase cell populations (C4, C5, C7) after EBV infection (**Figure 2A**), with decreased numbers of CD8^+^ T cells expressing CD45RA and CCR7, and increased representation of KLRG1, CD45RO, and CD103 positive CD8^+^ T cells (**Figure 2B**). PD-1 and CD69 molecules were also expressed by these populations (**Figure 2C**). These data confirmed that CD8^+^ T_EM_ lymphocytes expressing CD103 were the most abundant activated cells after EBV infection, whether they expressed CD69, or not (**Figure 2D, Supplementary Figure 3B**). A subset of CD4 T lymphocytes from IgG+ individuals also increased CD103 expression after EBV infection, but to a much lesser extent than CD8 T lymphocytes (**Supplementary Figure 3C,D**).

**Figure 2.**
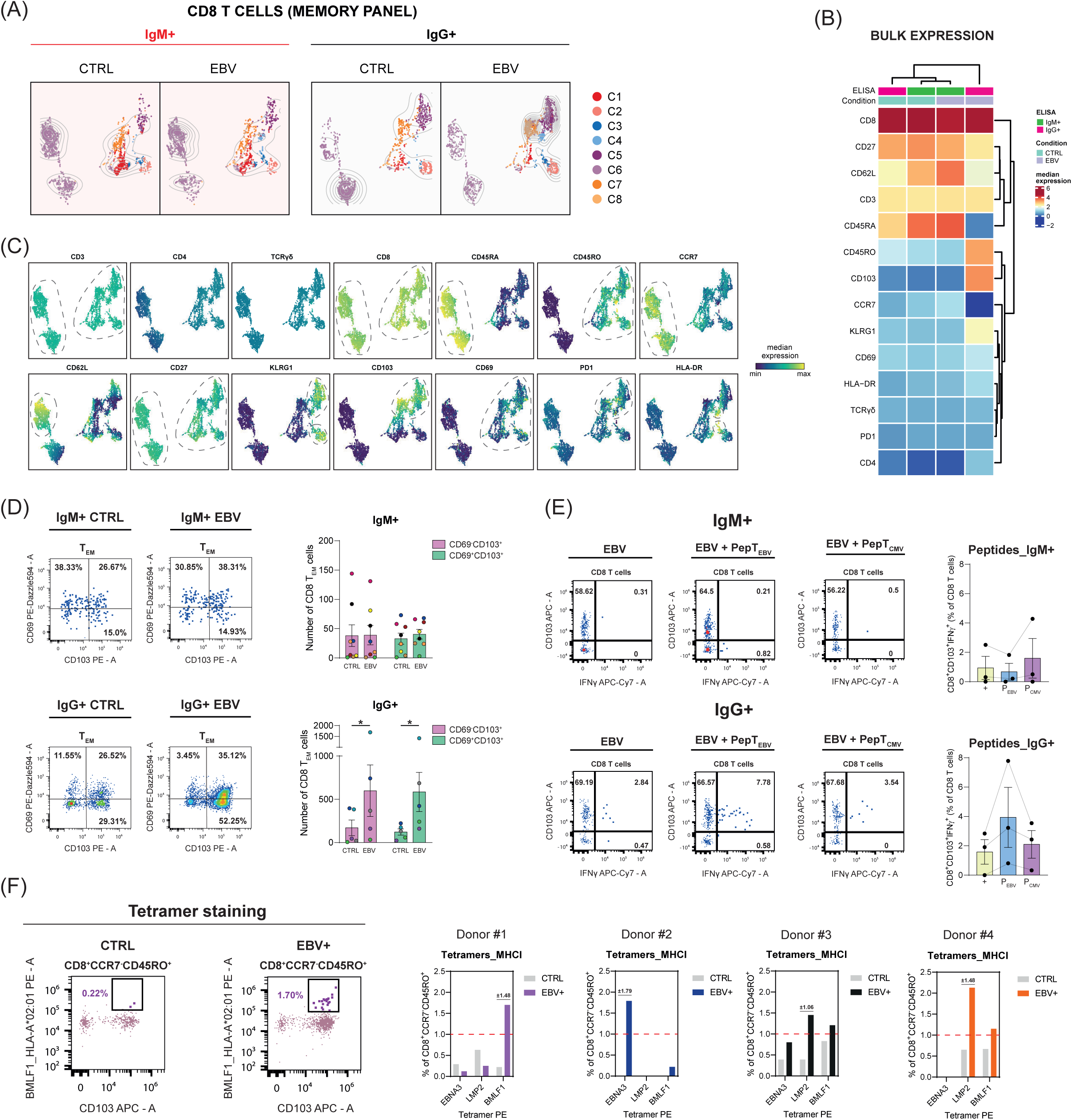
CD103^+^ effector memory CD8 T cells are enriched in IgG+ tonsils after infection with EBV. (A) UMAP analysing the CD8 T cells from the T memory panel split by condition and serology. Density contours were added to better illustrate the clusters that emerged after infection. (B) Heatmap representing the bulk expresion of markers per condition and serology for the flow cytometry T memory panel. (C) Feature-plots showing the expression of each marker from the T memory panel within CD8 T cells. (D) Representative flow cytometry dot-plots and quantification of CD103^+^ CD8 T cell populations in IgM+ and IgG+ samples after infection with EBV. Paired sample t-test (*<0.05). (E) Flow cytometry dot-plots and quantification of CD103^+^IFNγ^+^ CD8 T cells in IgM+ and IgG+ samples after infection and restimulation with EBV or HCMV peptides. (F) Determination of MHCI-tetramer binding to CD8^+^CCR7^-^CD45RO^+^ cells in infected IgG+ samples.

To test whether the activated CD103-expressing CD8 T lymphocytes in the tonsils contained EBV-antigen specific T cells, restimulation experiments, where the expression of IFN-γ and TNF-α was assessed after exposure to mixes of peptide epitopes from either EBV or HCMV, were carried out. CD103^+^ CD8 T lymphocytes from IgG+ individuals produced IFN-γ and/or TNF-α after restimulation with peptides from EBV, but not HCMV (**Figure 2E, Supplementary Figure 3E**). In parallel, HLA tetramers loaded with peptide epitopes from EBV specifically bound CD103^+^ CD8 T_EM_ cells from IgG+ individuals following infection with the virus, further confirming that the activated CD8^+^ T_RM_ cells contained T cells specific for EBV (**Figure 2F, Supplementary Figure 3F-H**).

### 3. Large amounts of IL-10 are detected after EBV-infection of tonsil cultures from IgM+ individuals

In order to define factors that might modulate the immune response to primary EBV infection, we used Luminex technology to quantify the levels of multiple cytokines and chemokines in the tonsil cultures. Supernatants from IgM+ and IgG+ tonsillar cultures as well as cultures of PBMCs from healthy EBV seropositive donors were compared in these experiments (**Supplementary Figure 4A**). The concentrations of cytokines were determined using logistic regressions, and the differences between control and infected conditions were quantified for each group of samples (**Figure 3A, Supplementary Figure 4B,C**). After EBV infection, the secretion of cytokines such as IL-10, MIP-1α, MIP-1β, and IL-27 was markedly elevated in tonsil cultures from IgM+ individuals. In contrast, little variation in the levels of these soluble factors was noted for IgG+ individuals (**Figure 3B,C**). Principal component analysis (PCA) of these data revealed that the patterns of cytokine production in EBV-infected IgM+ samples were clearly distinct from the other samples and that IL-10, MIP-1α, and IL-27 were important drivers of this separation. PBMC samples localised to another area of the graph with lower contribution of BCA-1, and showed higher expression of IFN-γ, IL-15, and MIG after infection (**Figure 3D**).

**Figure 3.**
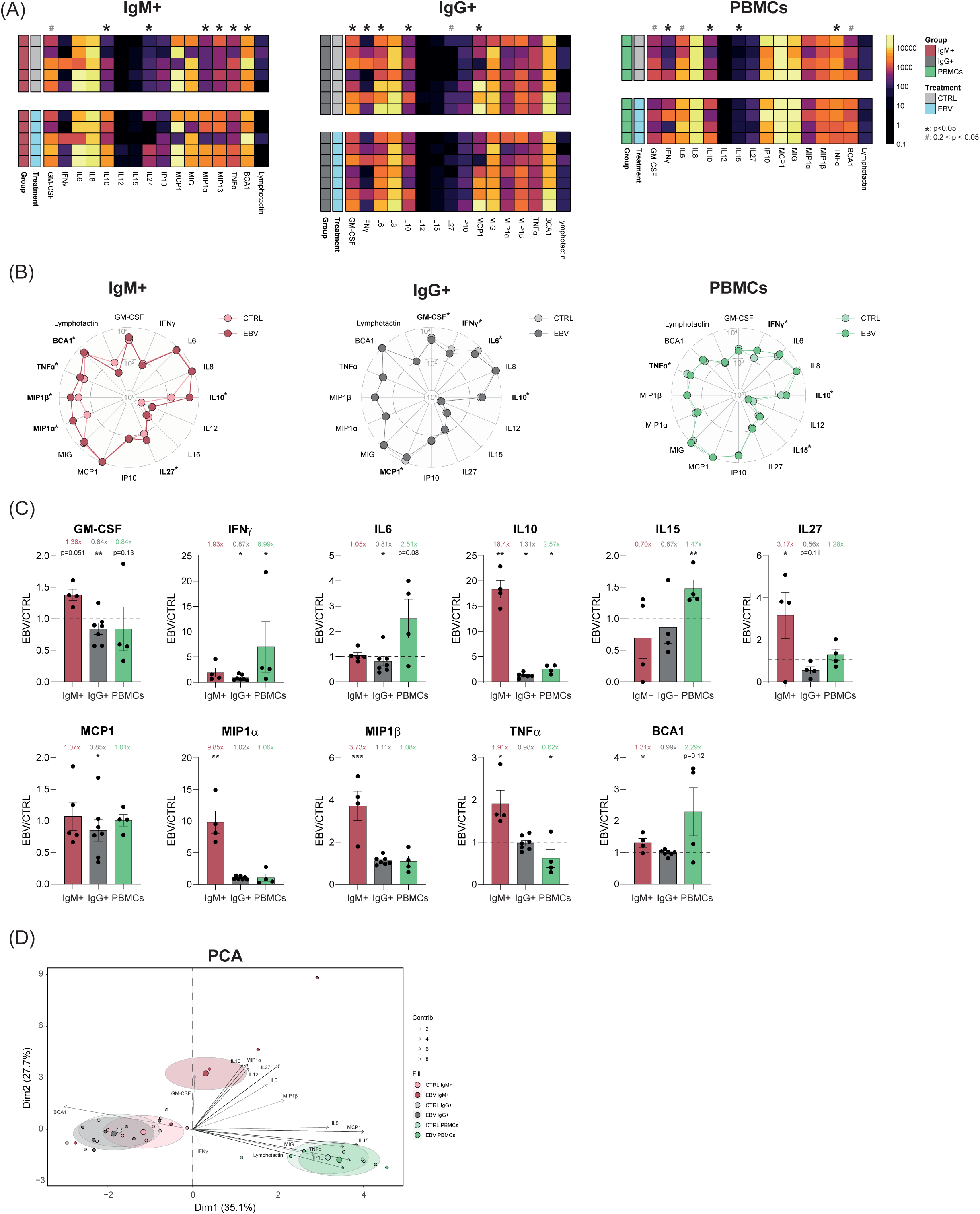
Cytokine determination. (A) Heatmap of raw values of cytokine measurements in a logarithmic scale for IgM+, IgG+, and PBMC samples, discriminating between control and infected conditions. Paired sample t-test (#<0.2, *<0.05). (B) Radar plots representing the mean of raw values for each cytokine between control and infected conditions in a logarithmic scale. Paired sample t-test (*p<0.05). (C) Bar-plots of the cytokine detection ratio between infected and control conditions. Only cytokines that were significant for paired sample t-test comparisons between control and infected groups were represented (*p<0.05, **p<0.01, ***p<0.001). Samples with abnormal values according to one-class SVM algorithm were excluded in each case. (D) PCA biplot discriminating per group and condition.

### 4. The EBV encoded IL-10 homologue negatively regulates T cell expansion in primary EBV infection of tonsils

IL-10 expressing cells were detected among the lymphoblastoid cells expanded after EBV infection in IgM+ tonsils (**Figure 4A**), but since the IL-10 specific monoclonal antibody (mAb) used in these experiments reacts with both human and EBV-encoded IL-10^17^, it was not clear whether the IL-10 being detected corresponded to the human cytokine or the viral IL-10 encoded by the EBV *BCRF1* gene. Although the amino acid identity between human and EBV IL-10 is very high, the nucleotide sequences show less similarity, thus oligonucleotide pairs that specifically amplified either human or viral IL-10 were optimised (**Supplementary Figure 5A,B**). RT-qPCR revealed that it was viral, not human, IL-10, that was expressed in infected IgM+ samples, especially on day 10 (**Figure 4B,C, Supplementary Figure 5C**).

**Figure 4.**
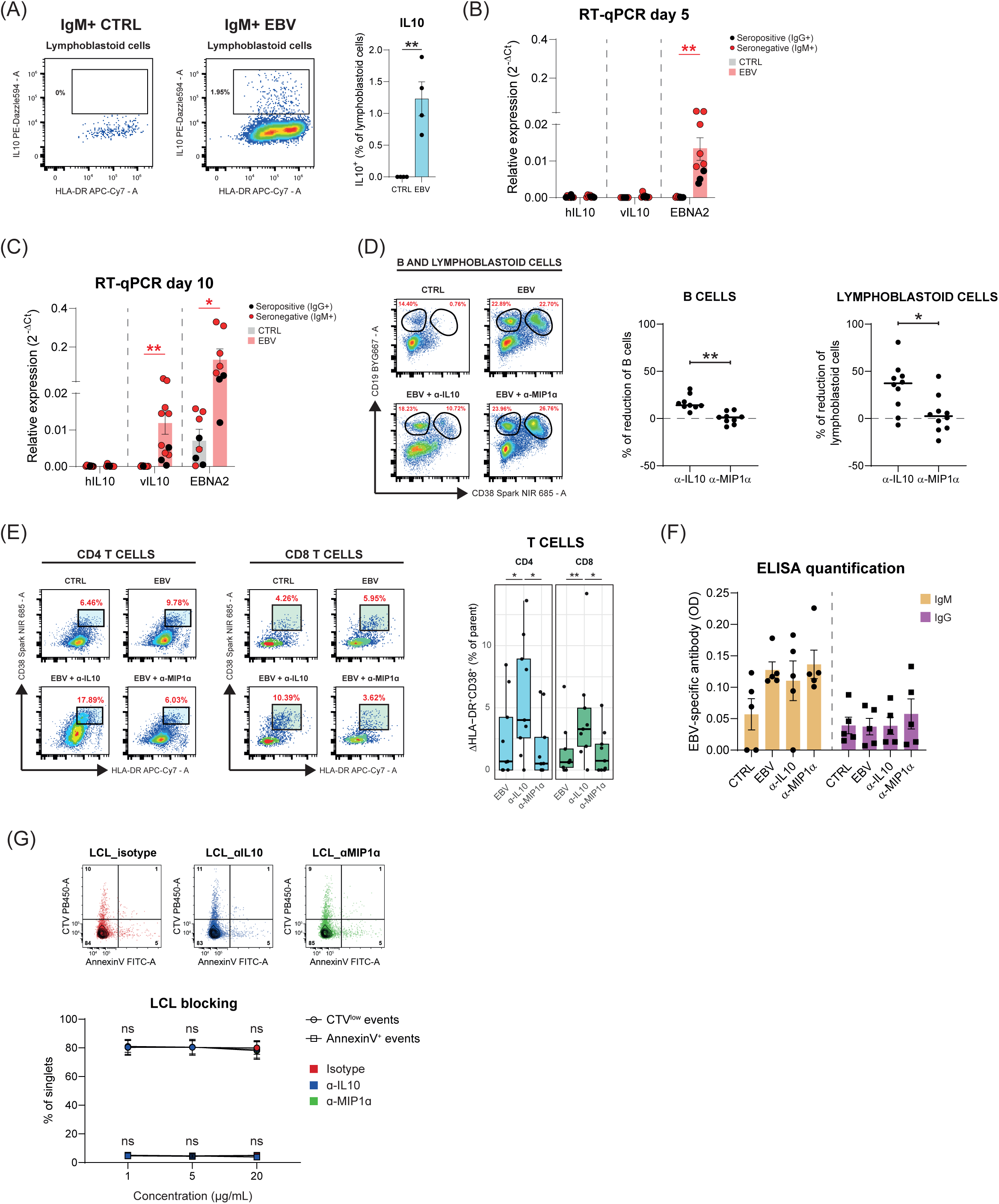
vIL-10 suppresses the immune response in IgM+ individuals. (A) IL-10 detection in the fraction of lymphoblastoid cells enriched after infection with EBV in IgM+ individuals after 10 days. (B) RT-qPCR relativised expression of hIL-10, vIL-10, and EBNA2 in control and infected conditions at day 5. (C) RT-qPCR relativised expression of hIL-10, vIL-10, and EBNA2 in control and infected conditions at day 10. (D) Percentage of reduction of B lymphocytes and lymphoblastoid cells in the infected + blocked conditions compared to the infected-only condition. Paired sample t-test (*<0.05, **<0.01). (E) Increase in the detection of HLA-DR and CD38 in CD4 and CD8 T lymphocytes after infection, infection + α-IL-10, and infection + α-MIP-1α. Paired sample t-test (*<0.05, **<0.01). (F) ELISAs quantifying IgM and IgG titers against EBV in the supernatant of tonsil organoids in control, infected, infected + α-IL-10, and infected + α-MIP-1α conditions. (G) Representative examples of CTV and AnnexinV staining of LCLs at day 6 after blocking with 20 µg/mL of α-IL-10 or α-MIP-1α and quantification of AnnexinV^+^ or CTV^low^ events per concentration and condition (n=3).

Since vIL-10 can exert anti-inflammatory effects similar to human IL-10^18,19,20^, a possible role for vIL-10 in modulating T lymphocyte activation in infected IgM+ tonsil samples was investigated by monoclonal antibody mediated neutralisation of IL-10. As a control, mAb blockade of the MIP-1α chemokine, that was also highly upregulated following infection of IgM+ tonsils, was carried out in parallel (**Figure 3**). Interestingly, blocking of IL-10, but not MIP-1α, led to a marked decrease in the outgrowth of lymphoblastoid cells and B lymphocytes (**Figure 4D**). This effect was accompanied by an increased detection of activated T lymphocytes (HLA-DR^+^CD38^+^) in the IL-10 blocking condition, consistent with an enhanced T cell immune response (**Figure 4E**). In proportion and number, the greatest increase in activation occurred in the CD4 compartment, but increased CD8 T-cell activation was also noted. IL-10 blockade did not lead to significant changes in the amounts of EBV-specific antibody produced in the culture or the Ig class of these antibodies (**Figure 4F**).

Culture of EBV-transformed lymphoblastoid cell lines (LCLs) with different doses of either IL-10 or MIP-1α specific blocking mAbs did not lead to significant changes in either LCL proliferation or death (**Figure 4G, Supplementary Figure 5D**), suggesting that the growth of EBV-transformed cells was not dependent on viral IL-10. Thus, these data are consistent with the hypothesis that the reduction in outgrowth of B lymphoblastoid cells after blocking IL-10 in the tonsil cultures was due to the increased activity of effector lymphocytes.

## DISCUSSION

Very little is known about how EBV infection progresses through natural routes in its native host, but as the virus is mainly transmitted from person to person in oral secretions, it is generally thought that it infects its main host, the human B cell, in lymphoid tissues like tonsils that are found in the oropharynx^1^. We have used EBV infection of tonsil organoid cultures as a model to study the initial interactions between the virus and the host immune response. Moreover, since in these experiments IgG antibodies to EBV were only detected in around one third of the tonsil donors, we had the opportunity to compare primary and memory immune responses to the virus in these cultures. Since only low levels of virus-neutralising antibodies were detected in these cultures, lymphocyte cytotoxicity was a strong candidate mechanism to play a key role in controlling the infection during the secondary response, and a marked activation of effector memory CD8 T lymphocytes expressing the CD103 integrin was noted in the tonsil organoid cultures^11^. The expression of this integrin defines tissue resident memory T lymphocytes and these CD8 T cells also upregulated expression of terminal differentiation markers such as KLRG1 and GZMB, showing a clear effector pattern, accompanied by notable expression of CD27, although in this case this marker seems to be constitutively expressed in tonsillar T cells^21^. The EBV specificity and activity of these lymphocytes were confirmed in experiments where the expanded CD8 T cells that expressed IFN-γ after restimulation with pools of peptide epitopes from EBV were found within the CD103^+^ population. Further, these memory CD8^+^CD103^+^ T cells bound HLA-I tetramers loaded with EBV epitopes coming from both lytic and latent expressed proteins. The detection of highly activated EBV-specific T-cells without marked expansion of total CD8^+^ numbers is reminiscent of observations made in the blood of children and young adults undergoing asymptomatic primary EBV infection, where a virus-specific CD8 T-cell response is able to control the infection without the exaggerated immune response associated with infectious mononucleosis^22,23^.

Vδ1 γδ T lymphocytes and NK cells, two other lymphocyte populations that have been shown to be able to recognise and kill EBV-infected cells, also increased during the secondary response^16^. Vδ1 cells upregulated markers of functional activity such as NKp30 and GZMB after infection, indicating that they likely contribute to eliminating EBV-infected cells. Interestingly, recent studies suggest that the numbers of Vδ1 gamma-delta T cells, that typically reside in tissues like skin, lungs, or intestine, are enriched in these sites in childhood, but gradually decline with age to the lower levels seen in adults^24^. It is tempting to speculate that this might be relevant to understanding why EBV infection in early life is often asymptomatic^25^, and that the likelihood of severe disease increases if primary infection occurs in adolescence. It is interesting that although most tonsil NK cells exhibit a CD56^bright^ phenotype, the strongest evidence of activation following EBV infection was found for CD56^dim^ cells with relatively low KIR expression. These lymphocytes may be related to the CD56^dim^ NKG2A^+^ immunoglobulin-like receptor^-^ NK cell subset that preferentially degranulates and proliferates on exposure to EBV-infected B cells expressing lytic antigens^26^.

In contrast, the cytotoxic cellular response was much more limited in tonsils undergoing primary infection, resulting in the outgrowth of B lymphoblastoid cells, without, at least in the time frame of these experiments, marked activation of cytotoxic lymphocytes.

While trying to understand the difference in immune control between EBV-infection of tonsil tissues from primary and virus-experienced donors, we noted that markedly increased levels of several soluble factors, including MIP-1α, MIP-1β, IL-27, and especially IL-10, were specific features of primary infection. RT-qPCR analysis showed that this IL-10 was actually of viral origin; the IL-10 homologue encoded in the EBV genome by the *BCRF1* gene that has marked immunomodulatory properties^27,28,29,30^. Although *BCRF1* has been characterised as a "late" viral gene expressed during the lytic phase of virus replication^31^, freshly EBV-infected human B cells have been shown to express vIL-10 within 4-6 h of infection and *in vitro* this continues over at least 28 days^18,32^. Indeed, it has been shown that mRNA able to drive transcription of *BCRF1* can even be found within infectious viral particles^33^.

Functionally, monoclonal antibody blockade of the viral IL-10 led to a reduced outgrowth of lymphoblastoid cells after primary infection. This effect correlated with enhanced CD4 and CD8 T lymphocyte responses, perhaps due to a polarisation toward Th1 that was previously inhibited. As neutralising IL-10 did not trigger any detectable increase in antibody production or an isotype switch towards IgG secretion, these observations suggest that stronger cell-mediated immunity is the key factor involved in better control of the growth of infected cells. This conclusion is consistent with multiple studies on immunity against EBV showing that CD8, but also CD4 T effector cells, are key responses that limit the growth of LCLs^34,35^. Further, our data showing that IL-10 blockade had no effect on the growth or survival of LCLs *in vitro* are consistent with studies showing that deletion of the viral IL-10 gene (*BCRF1*) from the EBV genome had no effect on EBV latent infection, B-lymphocyte proliferation into long-term lymphoblastoid cell lines, or replication^29,30^. These observations reinforce the suggestion that the mechanism underlying the restoration of immune control of EBV infection is enhanced T cell activation after IL-10 neutralisation.

Since, in most cases, EBV infection in childhood is asymptomatic, it seems likely that with time, *in vivo*, a cellular immune response will develop and control viral replication. However, it is plausible that this IL-10 dependent delayed activation of effective cytotoxic lymphocyte responders gives EBV a “window of opportunity” to establish a persistent, latent, infection and disseminate from the oropharynx to other sites in the body. Our model would be that while IL-10 inhibits local immunity in the tonsil, this cytokine is diluted in lymphatic vessels. In consequence, in draining lymph nodes, cross-presentation of debris from lytically infected cells as well as direct recognition of latently infected B cells (that will no longer express BCRF1) initiates the development of an effective cellular immune response that eventually seeds tissues with the EBV-specific cytotoxic lymphocytes that control EBV replication. This hypothesis is consistent with prior data from Dunmire *et al*^23^, who concluded that B cells are likely the major cell type to be first infected *in vivo* and showed that in many subjects EBV genomes could be detected in peripheral blood samples before detection in the oral cavity, again suggesting that migration of latently infected B cells is a major route for systemic dissemination of EBV infection.

Finally, it is interesting to note that whereas in our model of primary infection of tonsils IL-10 suppresses the immune response, elevated levels of IL-10 are a consistent feature of EBV-associated hemophagocytic lymphohistiocytosis (HLH), where the pathology is caused by systemic hyperproliferation of EBV-specific T lymphocytes^36,37,38,39,40^. Elevated serum IL-10 has also been reported to be a feature of post-transplant lymphoproliferative disorder (PTLD), infectious mononucleosis, and chronic active EBV (CAEBV) disease^41,42,43^. In most of these studies, the origin of the IL-10 (viral or human) in serum was not determined, however viral IL-10 was detected in around 20% of patients with infectious mononucleosis and almost one third of CAEBV patients, and viral IL-10 accounted for all the IL-10 detected in most or all of these samples^42,43^. Interestingly, in the CAEBV study, patients with severe disease displayed a trend toward higher mean viral IL-10 levels compared to patients with moderate disease. At first glance these data are surprising since IL-10 is generally thought of as an anti-inflammatory cytokine with immunosuppressive effects^44^. However, in some contexts, particularly in cancer, IL-10 administration has been reported to promote CD8 T cell activation associated with increased CD8 T cell infiltration in tissue, induced IFN-γ production, and favouring effective T cell memory responses^45,46,47,48^. Indeed, treatment with pegylated IL-10 has been shown to induce CD8 T cell immunity in human patients with solid tumors^49^. In HLH, Tang *et al*^38^ showed that high levels of circulating IL-10 and IL-18 in mice doubly transgenic for these cytokines triggered a lethal hyperinflammatory disease and that monoclonal antibody blockade of IL-10 protected against lethal disease in this HLH mouse model. Thus, it seems that depending on context, IL-10 cytokines can mediate either pro or anti-inflammatory effects. The basis of this divergence in outcome is not clear, but might be related to local, as opposed to systemic effects of the cytokine^50^. In either case, it seems reasonable to propose that the use of neutralising IL-10-specific antibodies, reactive with both cellular and viral IL-10, might be a useful adjunct to therapy in cases of EBV-associated HLH.

## MATERIALS AND METHODS

### 1. Samples

Whole tonsils from individuals undergoing surgery for obstructive sleep apnea or hypertrophy were collected in accordance with the Institutional Review Board (IRB) of the University Teaching Hospital “12 de Octubre” who determined that these samples had IRB Exempt Status, since the resected tonsil tissue is normally disposed of without further analysis. Written informed consent was obtained from the legal guardians of the children (aged 3-12 years) undergoing surgery for obstructive sleep apnea and/or hypertrophy, and overall, tonsil tissue was typically healthy. Whole tonsils were collected in saline after surgery and then immersed in an antimicrobial bath of Ham’s F12 medium (Gibco) containing Normocin (InvivoGen), penicillin, and streptomycin for 1 h at 4 °C for decontamination of the tissue. Tonsils were then briefly rinsed with PBS and processed as needed for culturing. Only de-identified demographic information was obtained.

### 2. Tonsil organoids

#### 2.1. Viruses

Aliquots of the B95-8 strain of EBV were obtained from the marmoset B-lymphoblastoid cell line (B95-8, ATCC CRL 1612). The cells were harvested in complete RPMI 1640 medium + 10% FBS, then centrifuged and plated in a 75 cm² flask containing complete RPMI 1640 + 5% FBS at a density of 10^6^ cells/mL for 96 hours. Subsequently, the cells were centrifuged at 2500 rpm for 10 minutes, and the supernatant containing EBV was collected using a 0.45 μm sterile filter. EBV-GFP was prepared from Akata cells by anti-Ig crosslinking^51^. Multiplicity of infection (MOI) was assessed by flow cytometry, after infection of tonsillar B cells with EBV-GFP for 48 hours. In other experiments, EBV-GFP was prepared by transfection of 293 cells containing the 2089 BAC clone^52^ (a gift from Dr. Obinna Chijioke, Institute of Experimental Immunology, University of Zurich, and Dr. Henri DeLeCleuse, German Cancer Research Center, Heidelberg) with plasmids that express the BZLF1 and BALF4 gene products (gifts from Dr. BE Gewurz, Brigham and Women’s Hospital, Boston) to produce EBV that is then titred by infection of Raji cells^53^.

#### 2.2. Transwell system

Tonsil samples were processed and cryopreserved as described by Wagar *et al*^15^. After thawing, cell debris was minimised by Ficoll density gradient separation, and isolated cells were resuspended at 6x10^7^ cells/mL in complete RPMI 1640 medium + 10% FBS. Cells were plated, 40 μL per well, onto permeable membranes (6.5 mm insert, 0.4-μm pore size), with the lower chamber consisting of 24-well plate sized wells covered with 450 μL of complete medium. In experimental conditions, 20 μL of EBV-enriched supernatant (MOI ∼0.01) was added directly to the cell-containing portion of the culture setup. Cultures were incubated at 37°C with 5% CO_2_ and humidity, and additional medium was added to the lower wells as needed.

#### 2.3. ELISA

For antigen coating, EBV and HSV-1 virions were purified from B95.8 cells and HSV1-infected Vero cells, respectively^54^. 100 μL of EBV or HSV preparations diluted to 0.5 μg/mL in Borate-buffered saline (10 mM Borate, 150 mM NaCl, pH 8.3) were added to each well of 96-well Maxisorp Nunc-Immuno plates and incubated overnight at 4°C. After incubation, the coating solutions were aspirated, and the ELISA plates were washed three times with 200 μL of PBS containing 0.05% Tween 20 (PBS-T) and subsequently dried. The plates were then blocked with PBS-casein (1× PBS blocker; Bio-Rad) for 1 hour at room temperature. Following this, 100 μL of culture supernatant diluted 1:2 or 1:5 in PBS-casein was added and plates were incubated overnight at 4°C. After three washes, 100 μL per well of the indicated detection antibodies (AffiniPure Rabbit Anti-Human IgM, Fcμ fragment specific; AffiniPure Rabbit Anti-Human IgG, Fcγ fragment specific from Jackson Labs) was added for 1 hour at room temperature. The plates were washed four times with PBS-T and then incubated in the dark at room temperature with 100 μL of Substrate Solution (OPD, Sigma-Aldrich) for approximately 3 minutes. Finally, 50 μL of stop solution (3 M H_2_SO_4_) was added to each well, and the optical density (OD) at 492 nm was measured using a microplate reader.

#### 2.4. Assay of virus neutralising antibodies

Samples of human serum or supernatants from tonsil cultures were incubated with medium containing 20,000 Green Raji Units (GRUs) of EBV-GFP for 30 minutes and then used to infect 100,000 Raji cells. After three days, the fraction of Raji cells expressing GFP was determined by flow cytometry.

#### 2.5. *In vitro* assays

Peptide stimulation was performed by incubating tonsil organoids, at day 10 after EBV infection, with mixes of known peptide epitopes from either EBV or HCMV (Miltenyi Biotec) followed by incubation for 6 hours in the presence of Brefeldin A (BioLegend, 5 μg/mL).

The IL-10 or MIP-1α blocking experiment was performed by adding 1 μg/mL of either α-IL-10 (clone JES3-9D7, BioLegend) or α-MIP-1α (clone #93321, R&D Systems) blocking antibodies to the culture immediately prior to tonsil infection with EBV. Antibody was replenished on day 6, and the cytometry analysis was done on day 10.

PBMC infection was performed after staining the PBMCs with CTV (ThermoFisher) and infecting them with EBV (MOI ∼5) for 2 hours at 37°C. After that, cells were washed 3 times in RPMI and plated in RPMI 1640 medium + 10% FBS at 2x10^5^ cells per well. Cultures were analysed on day 6.

Lymphoblastoid cell lines (LCLs) were generated by incubating PBMCs with PHA (5 μg/mL) for 30 minutes, followed by exposure to EBV (MOI∼ 5) for 2 hours in the presence of FK-506 (10 ng/mL). This was done in 96-well U-bottom plates at high confluence (2×10^5^ cells/well) in RPMI 1640 medium + 10% FBS. The medium was replaced weekly. As the cultures grew, LCLs were transferred to 24 and 6 well plates, and ultimately to 25 cm² flasks, over a period of approximately 28 days.

### 3. Flow cytometry

A complete list of the labelled anti-human antibodies utilised in this study is provided in **Supplementary Table 1**. For surface staining, cells were washed with PBS supplemented with 1% FBS, 0.5% BSA, and 0.05% sodium azide (PBA) before being incubated with the indicated antibodies for 30 minutes at 4°C. For tetramer staining, MHCI tetramers were stained at 4°C for 1 hour in darkness using a standard 1:100 dilution, and MHCII tetramers were stained at 37°C for 1 hour in darkness, using a concentration of 6 μg/mL. For intracellular staining, the surface-stained cells were washed in PBA, then fixed and permeabilised using either the True-Nuclear Transcription Factor Buffer Set (BioLegend) or PBA containing 4% paraformaldehyde and PBA containing 0.5% saponin. After this, antibodies were incubated with the cells for one hour at room temperature. Samples were then acquired on either a CytoFLEX (Beckman Coulter) or an Aurora 5L (Cytek) and analysed with CytExpert v2.5, SpectroFlo v2.2.0.3, or R software. For multiparametric flow cytometry analysis, FCS files exported from manual gating were processed using the R CATALYST package (v1.28.0), applying cofactors ranging from 150 to 3000.

### 4. Luminex assays

After infection of PBMCs with EBV for 6 days, or the infection of tonsil organoids with EBV for 10 days, the cytokine-enriched media were collected and centrifuged at 550g for 10 minutes. The supernatants were then stored at -80°C until further analysis using Luminex. The samples were analysed with magnetic Luminex® screening assays (R&D Systems) and a Luminex® 100™ analyser (Qiagen), following the manufacturer’s instructions. The concentration of each cytokine was interpolated via logistic regression of standard curves.

### 5. RT-qPCR

Total RNA was extracted from tonsil cultures using the TRIzol Plus kit (ThermoFisher), following the manufacturer’s protocol. For cDNA synthesis, RNA was used in a reverse transcription reaction according to SuperScript III (ThermoFisher) recommendations. The cDNA was then stored at -20°C until further use.

RT-qPCR was performed using the QuantStudio 5 Real-Time PCR system (ThermoFisher) with SYBR Green chemistry. The primers were designed using the Primer-BLAST tool and their sequences are detailed in **Supplementary Table 2**.

### 6. Statistics

Statistical analyses and graphical representations were performed using RStudio v4.2.1 or GraphPad Prism v8.3.0. Paired or unpaired comparisons between groups were described in figure legends. If not specified, the graphs shown in the figures represent the mean value ± SEM.

## DATA AVAILABILITY

Additional data from this study are available from the corresponding author upon request.

## AUTHOR’S CONTRIBUTIONS

Conceptualisation: A.F.G.J., H.T.R. Experiments: A.F.G.J. Formal analysis: A.F.G.J., A.S.C. Material support: L.I.G.G., I.J.H., M.V.G. Supervision: H.T.R. Funding acquisition: H.T.R., M.V.G. Writing-review and editing: A.F.G.J., H.T.R.

## ACKNOWLEDGEMENTS

We thank the tonsil donors and their families for donating samples to complete this study and the Pediatric ENT-Team (HU 12 de Octubre) for processing and coordinating the delivery of these samples. We also thank the NIH Tetramer Core Facility (NIH Contract 75N93020D00005 and RRID:SCR_026557) for providing tetramers of HLA-A2, -A3, -A11 and -DQB1*0602 loaded with peptide epitopes from BMLF1, EBNA1, EBNA3A, and LMP2 and labelled with phycoerythrin.

## SUPPLEMENTARY LEGENDS

**Supplementary Figure 1.** Characterisation of tonsil organoids after infection with EBV. (A) Microscopy examples of tonsil organoids in control and EBV-infected conditions. (B) Proportion of cells in control and infected conditions at day 10. Paired sample t-test (*<0.05, **<0.01). (C) Amplified proportion of lymphoblastoid cells in control and infected conditions at day 10. Paired sample t-test (**<0.01). (D) Quantification of EBV-GFP^+^ events within lymphoblastoid cells after infection of IgM+ tonsils at day 10. Paired sample t-test (*<0.05). (E) Graph representing the age of tonsil donors and the distribution between IgM+ and IgG+ samples. (F) Flow cytometry panels. (G) UMAP of T cells split by condition and serology. 10,000 cells per donor were plotted. The most differential clusters (C5 and C9) were highlighted. (H) Representative flow cytometry example of a seropositive donor (IgG+) upregulating HLA-DR, CD38, KLRG1, and GZMB after infection with EBV. (I) Feature-plots highlighting the expression of some markers in T cells. (J) Quantification of cells in the cytotoxic CD4 T cell cluster (C6) in control and infected conditions at day 10. (K) Feature-plot and flow cytometry plot showing the detection of FOXP3 C9 in the flow cytometry T cell panel. (L) UMAP of Vδ1 | Vδ2 | MAIT split by condition and serology. All cells were plotted. The most differential cluster (C3) was highlighted. (M) UMAP of NK cells split by condition and serology. All cells were plotted. The most differential cluster (C2) was highlighted. (N) Quantification of cells in C1, C3, C4, and C5 in control and infected conditions at day 10. Paired sample t-test (*<0.05).

**Supplementary Figure 2.** Characterisation of tonsil organoids after infection with EBV (A) Gating strategy of ILCs. (B) Number and proportion of ILCs in control and infected conditions at day 5. (C) Ratio of HLA-DR expression in ILCs between the infected and control conditions. (D) Gating strategy of macrophages. (E) Number and proportion of macrophages in control and infected conditions at day 5. Paired sample t-test (*<0.05). (F) Ratio of HLA-DR or CD80 expression in macrophages between the infected and control conditions.

**Supplementary Figure 3.** (A) Flow cytometry T cell memory panel. (B) Representative flow cytometry dot-plots showing percentages of naive, effector memory, effector memory reexpressing CD45RA, and central memory CD8 T cell populations in IgM+ and IgG+ samples. Paired sample t-test (*<0.05). (C) UMAP analysing the CD4 T cells from the T memory panel split by condition and serology. 5,000 cells per donor were plotted. The most differential cluster (C3) was highlighted. (D) Feature-plots showing the expression of each marker from the T memory panel within CD4 T cells. CD103 and PD-1 were highlighted for C3. (E) Quantification of IFNγ^+^TNFα^+^ CD8 T cells in IgM+ and IgG+ samples after infection and restimulation with EBV or HCMV peptides. (F) List of tetramers specific for EBV antigens. (G) Examples of MHCI-tetramer binding to CD8^+^CCR7^-^CD45RO^+^ cells in infected IgG+ samples. (H) Example and quantification of MHCII-tetramer binding to CD4^+^CCR7^-^CD45RO^+^ cells in infected IgG+ samples.

**Supplementary Figure 4.** Luminex assays. (A) Selection of PBMCs for Luminex assays according to CTV diliution of CD4, CD8, or NK cells after infection with EBV. (B) Logistic regression of each cytokine used in Luminex assays. (C) Bar-plots of the cytokine values obtained for each group and condition, in a logarithmic scale. Samples with abnormal values according to one-class SVM algorithm were excluded in each case. Paired sample t-test (*p<0.05, **p<0.01, ***p<0.001).

**Supplementary Figure 5.** IL-10 determination. (A) Clustal-ω alignment between the amino acid sequences of BCRF1 and human IL-10. Perfect and conservative matches were highlighted. (B) Clustal-ω alignment between the nucleic acid sequences of *BCRF1* and human IL-10. Perfect matches were highlighted. The bold sequences represent the primers designed to discriminate human or viral IL-10 signal in qPCR. (C) RT-qPCR amplification plot of the marmoset B95-8 cDNA. Detection of viral IL-10, human IL-10, and GAPDH are represented. (D) Representative examples of propidium iodide and AnnexinV staining of LCLs at day 6 after blocking with 20 µg/mL of α-IL-10 or α-MIP-1α and quantification of AnnexinV^+^PI^+^ events per concentration and condition.

**Supplementary Table 1.** Flow cytometry reagents. Supplementary Table 2. RT-qPCR primers.

