## Supplementary data for "The primary and secondary immune response to Epstein-Barr virus infection of human tonsil organoids"

**SUPP. TABLE 1**

| | Fluorophore | Clone | Supplier |
|---|---|---|---|
| **Viability** | | | |
| AnnexinV | FITC | - | Thermo Fisher |
| Propidium iodide | - | - | Thermo Fisher |
| LIVE/DEAD Fixable Blue | - | - | Thermo Fisher |
| **Surface** | | | |
| c-kit | PE-Dazzle594 | 104D2 | BioLegend |
| CCR5 | Pacific Blue | HEK/1/85a | BioLegend |
| CCR7 | PerCP5.5 | G043H7 | BioLegend |
| CD103 | PE | Ber-ACT8 | BioLegend |
| CD103 | APC | Ber-ACT8 | BioLegend |
| CD11c | APC | Bu15 | BioLegend |
| CD127 | PE-Cy7 | A019D5 | BioLegend |
| CD14 | PE | M5E2 | BioLegend |
| CD16 | PE-Cy7 | 3G8 | BioLegend |
| CD16 | V500 | 3G8 | BD Biosciences |
| CD161 | AF700 | HP-3G10 | BioLegend |
| CD19 | BYG667 | SJ25C1 | CYTEK |
| CD19 | FITC | HIB19 | BioLegend |
| CD25 | PE | BC96 | BioLegend |
| CD27 | FITC | M-T271 | BioLegend |
| CD3 | FITC | OKT3 | BioLegend |
| CD3 | Pacific Blue | UCHT1 | BioLegend |
| CD3 | PerCP5.5 | OKT3 | BioLegend |
| CD3 | BV605 | OKT3 | BioLegend |
| CD303 | FITC | 201A | BioLegend |
| CD34 | FITC | 581 | BioLegend |
| CD38 | Spark-NIR685 | HIT2 | BioLegend |
| CD4 | APC | OKT4 | BioLegend |
| CD4 | V500 | RPA-T4 | BD Biosciences |
| CD45RA | Pacific Blue | HI100 | BioLegend |
| CD45RO | PE-Cy5 | UCHL1 | BioLegend |
| CD56 | PE | MEM-188 | BioLegend |
| CD56 | PE-Cy5 | MEM-188 | BioLegend |
| CD57 | PE-Dazzle594 | HNK-1 | BioLegend |
| CD62L | APC-Cy7 | DREG-56 | BioLegend |
| CD69 | BV421 | FN50 | BioLegend |
| CD69 | PE-Dazzle594 | FN50 | BioLegend |
| CD8 | PE-Cy7 | SK1 | BioLegend |
| CD8 | Spark-Plus UV395 | RPA-T8 | BioLegend |
| CD80 | PE-Cy5 | 2D10 | BioLegend |
| CD94 | FITC | DX22 | BioLegend |
| CRTH2 | PE | BM16 | BioLegend |
| CTV | - | - | Thermo Fisher |
| FceRIa | FITC | AER-37 (CRA-1) | BioLegend |
| HLA-DR | APC-Cy7 | L243 | BioLegend |
| HLA-DR | Spark-Blue574 | Tü39 | BioLegend |
| KIR2DL1 | PE-Cy7 | HP-MA4 | BioLegend |
| KIR2DL2/L3 | PE-Cy7 | DX27 | BioLegend |
| KLRG1 | APC | SA231A2 | BioLegend |
| KLRG1 | PE-Cy7 | SA231A2 | BioLegend |
| NKG2D | PE | 1D11 | BioLegend |
| NKp30 | APC | P30-15 | BioLegend |
| NKp44 | APC | P44-8 | BioLegend |
| NKp80 | APC | 5D12 | BioLegend |
| PD-1 | APC | NAT105 | BioLegend |
| TCRγδ | AF660 | B1 | BioLegend |
| TCRVα7.2 | PE-Cy7 | 3C10 | BioLegend |
| TRDV1 | PE | TS8.2 | Thermo Fisher |
| TRDV2 | FITC | B6 | BioLegend |
| **Intracellular** | | | |
| FOXP3 | BV421 | 206D | BioLegend |
| GZMB | PE-Dazzle594 | QA18A28 | BioLegend |
| GZMB | PerCP5.5 | QA16A02 | BioLegend |
| IFNγ | BUV805 | 4S.B3 | Thermo Fisher |
| IFNγ | PE-Cy7 | 4S.B3 | BioLegend |
| IL10 | PE | JES3-19F1 | BioLegend |
| TNFα | PE | MAb11 | BioLegend |

### SUPP. TABLE 2

| Gene | Sequence |
|---|---|
| GAPDH (F) | CACCCACTCCTCCACCTTTGAC |
| GAPDH (R) | GTCCACCACCCTGTTGCTGTAG |
| BcRF1 (F) | GACCAATGTGACAATTTTCCCC |
| BcRF1 (R) | AAGAATTGACAATGGTCTTTGGC |
| hIL10 (F) | TCCCAGGCAACCTGCCTAAC |
| hIL10 (R) | GAGTTCACATGCGCCTTGATC |
| EBNA2 (F) | CCCATCCAATGCCGCCCCCG |
| EBNA2 (R) | GAGGTCTTTTACTGGGTCCC |

**SUPP. 1**

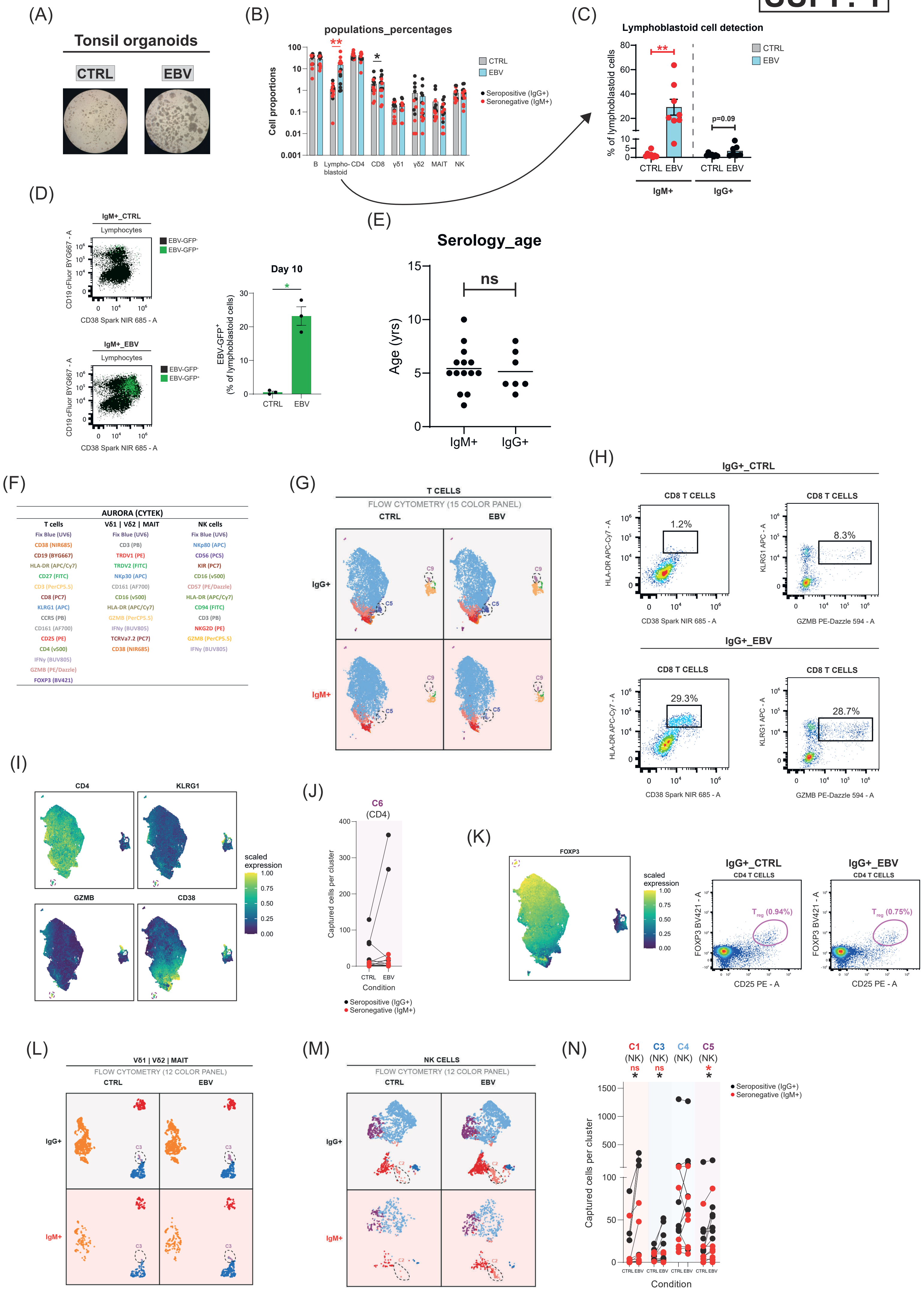
(A)
Tonsil organoids
CTRL
EBV
(B)
populations_percentages
Cell proportions
100
10
1
0.1
0.01
0.001
B
Lympho-blastoid
CD4
CD8
γδ1
γδ2
MAIT
NK
CTRL
EBV
Seropositive (IgG+)
Seronegative (IgM+)
(C)
Lymphoblastoid cell detection
% of lymphoblastoid cells
p=0.09
CTRL
EBV
IgM+
IgG+
(D)
IgM+_CTRL
Lymphocytes
EBV-GFP-
EBV-GFP+
CD19 cFluor BYG667 - A
CD38 Spark NIR 685 - A
IgM+_EBV
Day 10
EBV-GFP+ (% of lymphoblastoid cells)
(E)
Serology_age
ns
Age (yrs)
IgM+
IgG+
(F)
AURORA (CYTEK)
T cells
Vδ1 | Vδ2 | MAIT
NK cells
Fix Blue (UV6)
CD38 (NIR685)
CD19 (BYG667)
HLA-DR (APC/Cy7)
CD27 (FITC)
CD3 (PerCP5.5)
CD8 (PC7)
KLRG1 (APC)
CCR5 (PB)
CD161 (AF700)
CD25 (PE)
CD4 (v500)
IFNγ (BUV805)
GZMB (PE/Dazzle)
FOXP3 (BV421)
Fix Blue (UV6)
CD3 (PB)
TRDV1 (PE)
TRDV2 (FITC)
NKp30 (APC)
CD161 (AF700)
CD16 (v500)
HLA-DR (APC/Cy7)
GZMB (PerCP5.5)
IFNγ (BUV805)
TCRVa7.2 (PC7)
CD38 (NIR685)
Fix Blue (UV6)
NKp80 (APC)
CD56 (PC5)
KIR (PC7)
CD16 (v500)
CD57 (PE/Dazzle)
HLA-DR (APC/Cy7)
CD94 (FITC)
CD3 (PB)
NKG2D (PE)
GZMB (PerCP5.5)
IFNγ (BUV805)
(G)
T CELLS
FLOW CYTOMETRY (15 COLOR PANEL)
CTRL
EBV
IgG+
IgM+
C5
C9
(H)
IgG+_CTRL
CD8 T CELLS
1.2%
8.3%
HLA-DR APC-Cy7 - A
KLRG1 APC - A
CD38 Spark NIR 685 - A
GZMB PE-Dazzle 594 - A
IgG+_EBV
29.3%
28.7%
(I)
CD4
KLRG1
GZMB
CD38
scaled expression
1.00
0.75
0.50
0.25
0.00
(J)
C6
(CD4)
Captured cells per cluster
Condition
Seropositive (IgG+)
Seronegative (IgM+)
(K)
FOXP3
IgG+_CTRL
CD4 T CELLS
Treg (0.94%)
IgG+_EBV
Treg (0.75%)
FOXP3 BV421 - A
CD25 PE - A
(L)
Vδ1 | Vδ2 | MAIT
FLOW CYTOMETRY (12 COLOR PANEL)
C3
(M)
NK CELLS
FLOW CYTOMETRY (12 COLOR PANEL)
C2
(N)
C1 (NK)
C3 (NK)
C4 (NK)
C5 (NK)
ns
Captured cells per cluster
Condition

SUPP. 2

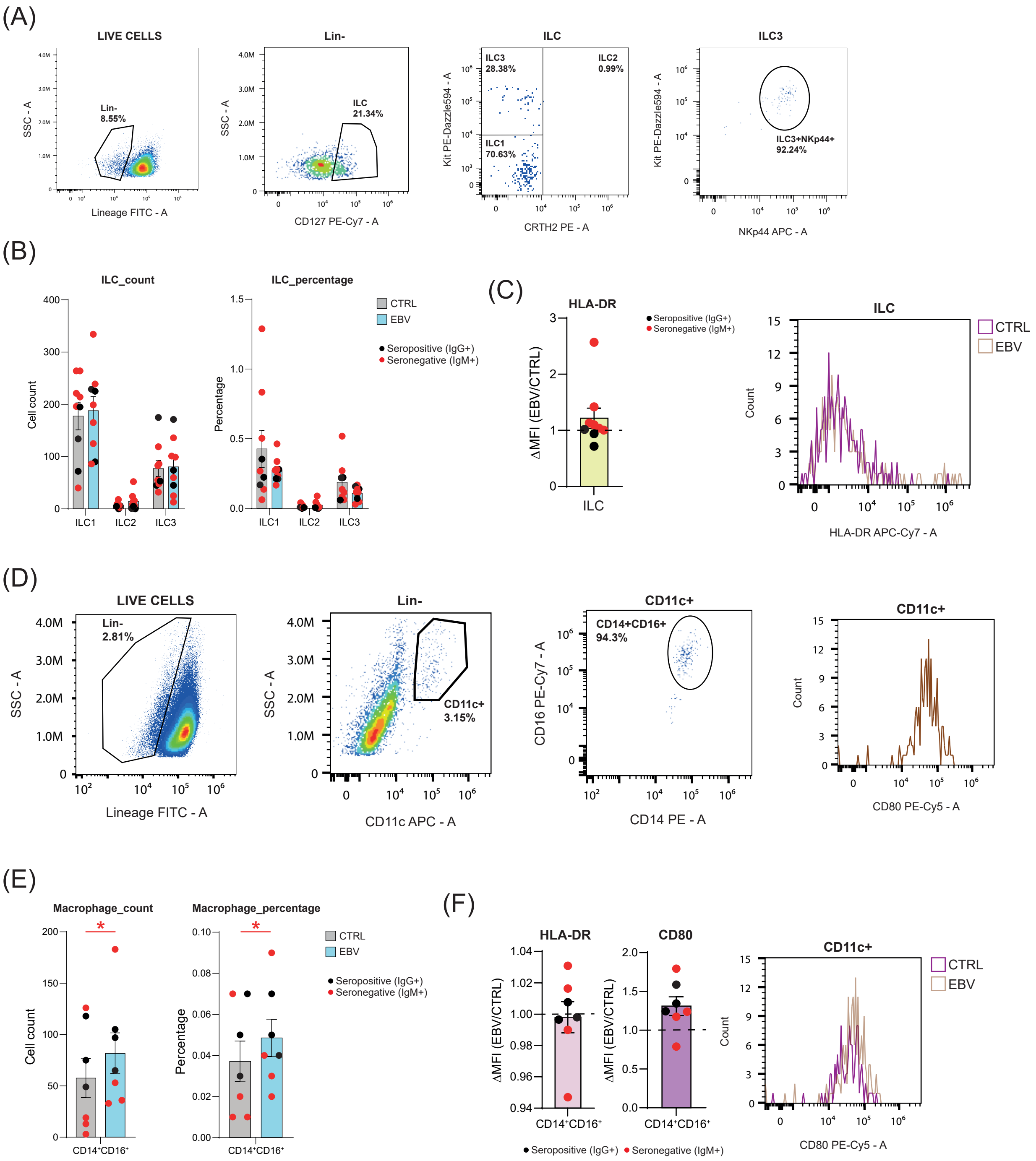

(A)
LIVE CELLS
Lin-
8.55%
SSC - A
Lineage FITC - A
Lin-
ILC
21.34%
CD127 PE-Cy7 - A
ILC
ILC3
28.38%
ILC2
0.99%
ILC1
70.63%
Kit PE-Dazzle594 - A
CRTH2 PE - A
ILC3
ILC3+NKp44+
92.24%
NKp44 APC - A
(B)
ILC_count
Cell count
ILC_percentage
Percentage
ILC1
ILC2
ILC3
CTRL
EBV
Seropositive (IgG+)
Seronegative (IgM+)
(C)
HLA-DR
ΔMFI (EBV/CTRL)
ILC
Seropositive (IgG+)
Seronegative (IgM+)
ILC
CTRL
EBV
Count
HLA-DR APC-Cy7 - A
(D)
LIVE CELLS
Lin-
2.81%
SSC - A
Lineage FITC - A
Lin-
CD11c+
3.15%
CD11c APC - A
CD11c+
CD14+CD16+
94.3%
CD16 PE-Cy7 - A
CD14 PE - A
CD11c+
Count
CD80 PE-Cy5 - A
(E)
Macrophage_count
Cell count
Macrophage_percentage
Percentage
CD14⁺CD16⁺
CTRL
EBV
Seropositive (IgG+)
Seronegative (IgM+)
(F)
HLA-DR
CD80
ΔMFI (EBV/CTRL)
CD14⁺CD16⁺
Seropositive (IgG+)
Seronegative (IgM+)
CD11c+
CTRL
EBV
Count
CD80 PE-Cy5 - A

### SUPP. 3

(A)

| T memory |
|---|
| Fix Blue (UV6) |
| CD3 (BV605) |
| CD4 (V500) |
| CD27 (FITC) |
| CD62L (APC/Cy7) |
| CD69 (PE/Dazzle) |
| TCRγδ (AF660) |
| PD1 (APC) |
| CD8 (SparkPlus UV395) |
| CD45RA (PB) |
| CD45RO (PC5) |
| CCR7 (PerCP5.5) |
| KLRG1 (PC7) |
| CD103 (PE) |
| HLA-DR (SparkBlue574) |

(B)

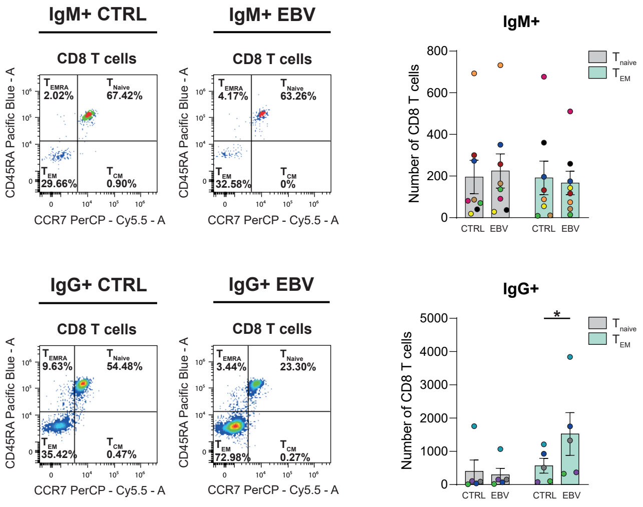


(C)

#### CD4 T CELLS (MEMORY PANEL)

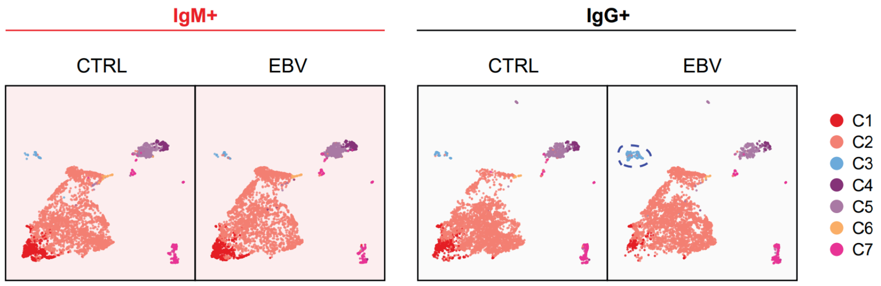


(D)

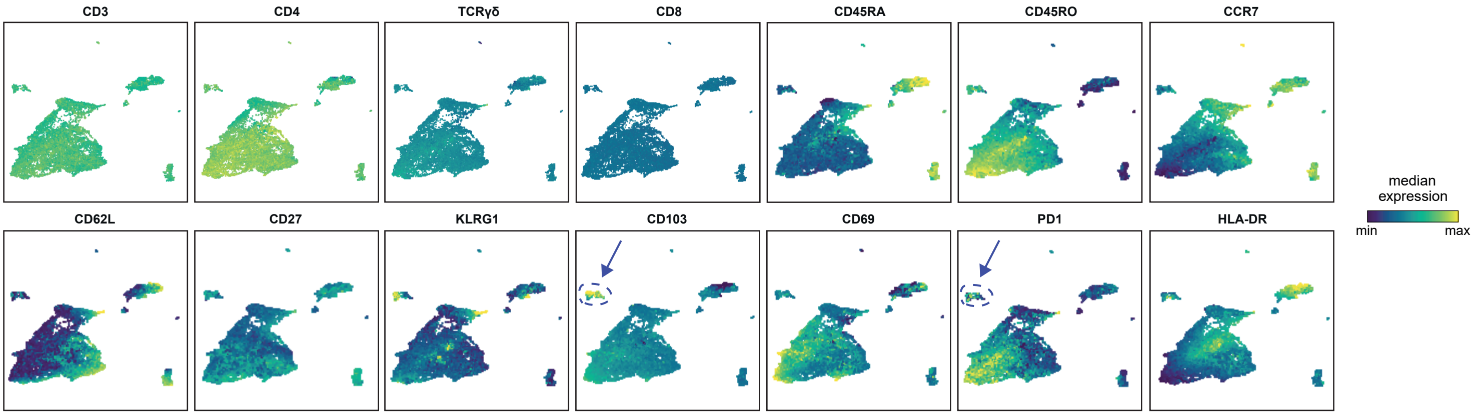


(E)

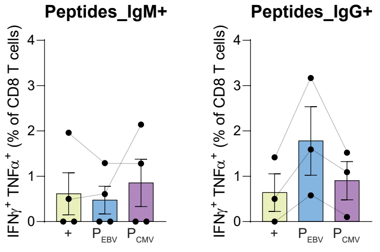


(F)

| Tetramer | Antigen | Restriction | MHC |
|---|---|---|---|
| RLRAEAQVK | EBNA3A | HLA-A*11:01 | I |
| RLRAEAQVK | EBNA3A | HLA-A*03:01 | I |
| CLGGLLTMV | LMP2 | HLA-A*02:01 | I |
| GLCTLVAML | BMLF1 | HLA-A*02:01 | I |
| RALLARSHVERTTD | EBNA1 | DQB1*06:02/DQA1*01:02 | II |
| PVSKMRMATPLLMQA | CLIP (ctrl) | DQB1*06:02/DQA1*01:02 | II |

(G)

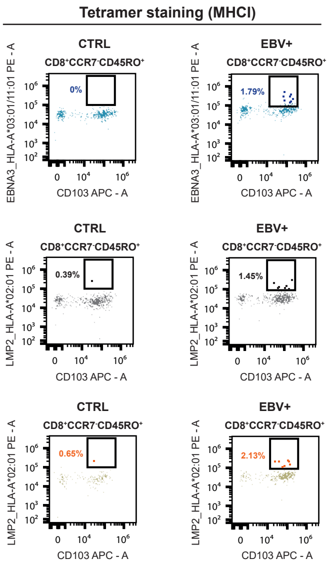


(H)

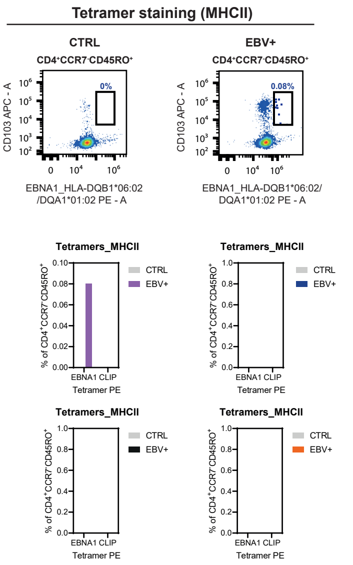

### SUPP. 4

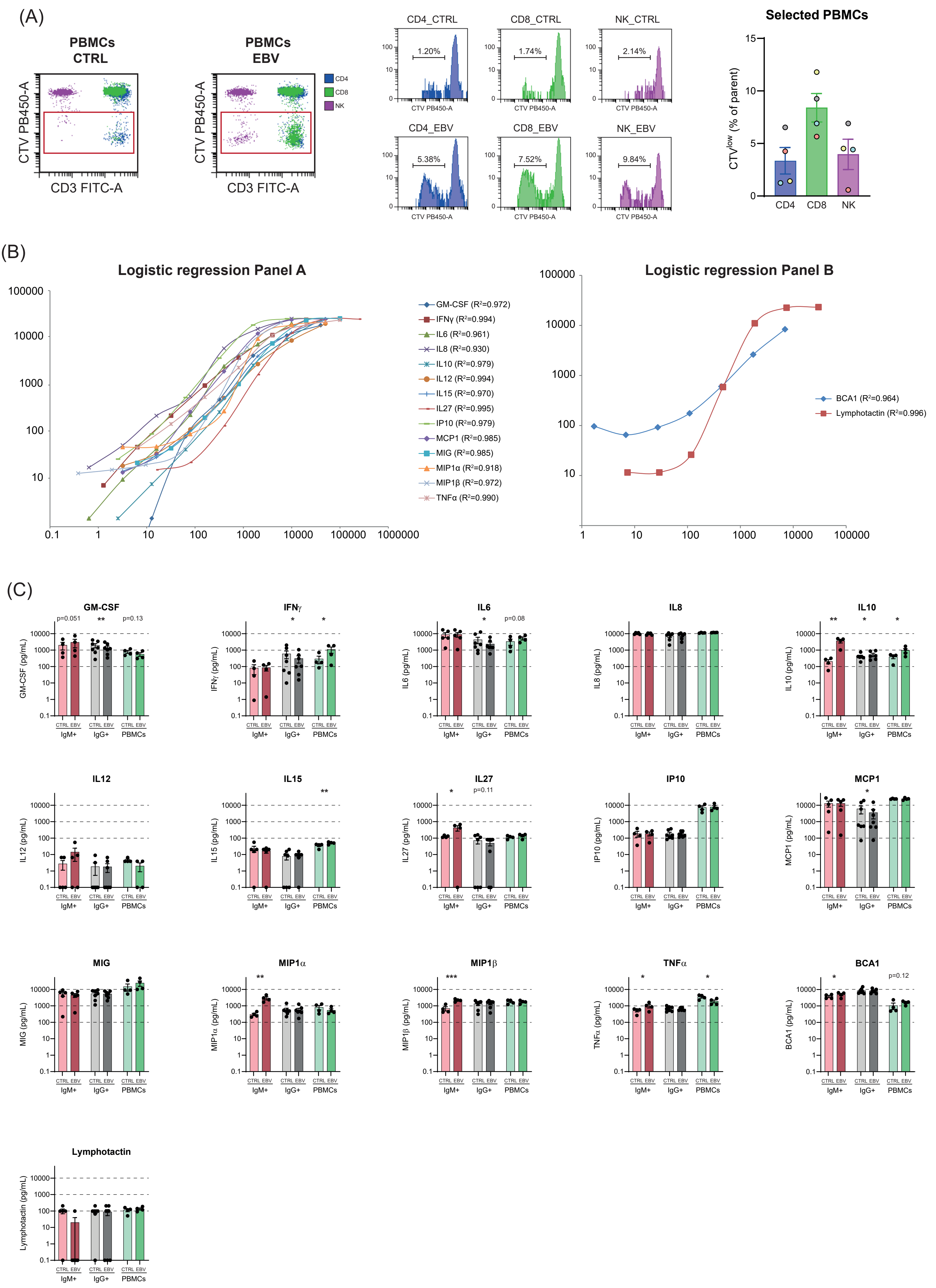
(A)
PBMCs CTRL
PBMCs EBV
CTV PB450-A
CD3 FITC-A
CD4
CD8
NK
CD4_CTRL
CD8_CTRL
NK_CTRL
1.20%
1.74%
2.14%
CD4_EBV
CD8_EBV
NK_EBV
5.38%
7.52%
9.84%
Selected PBMCs
CTVlow (% of parent)
(B)
Logistic regression Panel A
GM-CSF (R²=0.972)
IFNγ (R²=0.994)
IL6 (R²=0.961)
IL8 (R²=0.930)
IL10 (R²=0.979)
IL12 (R²=0.994)
IL15 (R²=0.970)
IL27 (R²=0.995)
IP10 (R²=0.979)
MCP1 (R²=0.985)
MIG (R²=0.985)
MIP1α (R²=0.918)
MIP1β (R²=0.972)
TNFα (R²=0.990)
Logistic regression Panel B
BCA1 (R²=0.964)
Lymphotactin (R²=0.996)
(C)
GM-CSF
IFNγ
IL6
IL8
IL10
IL12
IL15
IL27
IP10
MCP1
MIG
MIP1α
MIP1β
TNFα
BCA1
Lymphotactin
p=0.051
p=0.13
p=0.08
p=0.11
p=0.12
CTRL EBV
IgM+
IgG+
PBMCs

### SUPP. 5

(A)

```
BCRF1            MERRLVVTLQCLVLLYLA---PECG--GTDQ----CDNFPQMLRDLRDAFSRVKTFFQTK51
hIL-10           --MHSSALLCCLVLLTGVRASPGQGTQSENSCTHFPGNLPNMLRDLRDAFSRVKTFFQMK58
                    :  . * *****  .   *  *  . :.     .*:*:***************** *

BCRF1            DEVDNLLLKESLLEDFKGYLGCQALSEMIQFYLEEVMPQAENQDPEAKDHVNSLGENLKT111
hIL-10           DQLDNLLLKESLLEDFKGYLGCQALSEMIQFYLEEVMPQAENQDPDIKAHVNSLGENLKT118
                 *::******************************************: * ***********

BCRF1            LRLRLRRCHRFLPCENKSKAVEQIKNAFNKLQEKGIYKAMSEFDIFINYIEAYMTIKAR-170
hIL-10           LRLRLRRCHRFLPCENKSKAVEQVKNAFNKLQEKGIYKAMSEFDIFINYIEAYMTMKIRN178
                 ***********************:*******************************:* *
```

(B)

```
BCRF1           atggag------------cgaaggttagtggtcactctgcag----tgcctg-----gtg 39
hIL10           atgcacagctcagcactgctctgttgcctggtcctcctgactggggtgagggccagccca 60
                *** *             *   * *   *****   ***       **   *

BCRF1           ctgctttacctggcacctgagtgtggaggtacagaccaatgtgacaattttccccaaatg 99
hIL10           ggccagggcacccagtctgagaacagctgcacccacttcccaggcaacctgcctaacatg 120
                   *    *       *****    *  * **  **      * ***  * **  * ***

BCRF1           ttgagggacctaagagatgccttcagtcgtgttaaaacctttttccagacaaaggacgag 159
hIL10           cttcgagatctccgagatgccttcagcagagtgaagactttctttcaaatgaaggatcag 180
                 *  * ** **  *************  * ** ** ** ** ** ** *  *****  **

BCRF1           gtagataaccttttgctcaaggagtctctgctagaggactttaagggctaccttggatgc 219
hIL10           ctggacaacttgttgttaaaggagtccttgctggaggactttaagggttacctgggttgc 240
                 * ** *** * *** * ********  **** ************** ***** ** ***

BCRF1           caggccctgtcagaaatgatccaattctacctggaggaagtcatgccacaggctgaaaac 279
hIL10           caagccttgtctgagatgatccagttttacctggaggaggtgatgccccaagctgagaac 300
                ** *** **** ** ******** ** *********** ** ***** ** ***** ***

BCRF1           caggaccctgaagccaaagaccatgtcaattctttgggtgaaaatctaaagaccctacgg 339
hIL10           caagacccagacatcaaggcgcatgtgaactccctgggggagaacctgaagaccctcagg 360
                ** ***** **   *** *  ***** ** **  **** ** ** ********  **

BCRF1           ctccgcctgcgcaggtgccacaggttcctgccgtgtgagaacaagagtaaagctgtggaa 399
hIL10           ctgaggctacggcgctgtcatcgatttcttccctgtgaaaacaagagcaaggccgtggag 420
                **  * ** **  * ** **  * ** ** ** ***** ******** ** ** *****

BCRF1           cagataaaaaatgcctttaacaagctgcaggaaaaaggaatttacaaagccatgagtgaa 459
hIL10           caggtgaagaatgcctttaataagctccaagagaaaggcatctacaaagccatgagtgag 480
                *** * ** *********** ***** ** ** ***** ** *****************

BCRF1           tttgacatttttattaactacatagaagcatacatgacaattaaagccaggtga---    513
hIL10           tttgacatcttcatcaactacatagaagcctacatgacaatgaagatacgaaactga    537
                ******** ** ** ************** *********** **     *
```

(C)

B95-8 cell line

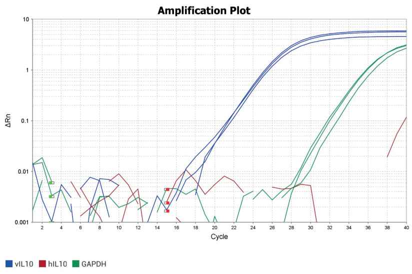


(D)

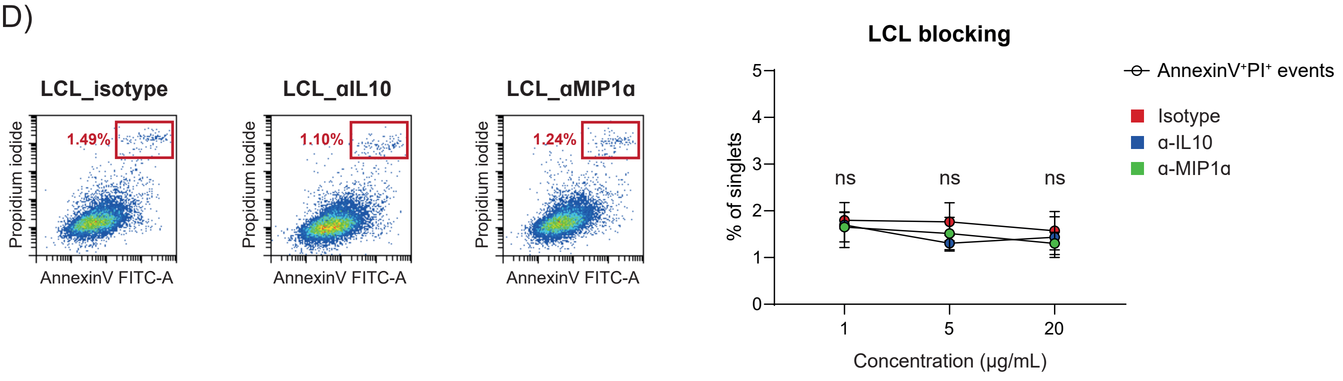
